# Physiological characterization in a somaclonal mutant for fruit shape derived from persimmon ‘Hiratanenashi’

**DOI:** 10.1101/2025.05.19.654774

**Authors:** Ayano Horiuchi, Ryusuke Matsuzaki, Noriyuki Onoue, Koichiro Ushijima, Yasutaka Kubo, Takashi Akagi, Mai F. Minamikawa

**Affiliations:** Graduate School of Environmental and Life Science, Okayama University, 1-1-1 Tsushima-naka, Okayama, 700-8530, Japan; Graduate School of Horticulture, Chiba University, 648 Matsudo, Matsudo, Chiba 271-8510, Japan; Institute of Fruit Tree and Tea Science, NARO, 301-2 Mitsu, Akitsu, Higashi-hiroshima, Hiroshima, 739-2494, Japan; JST, PRESTO, Kawaguchi-shi, Saitama, 332-0012, Japan; Institute for Advanced Academic Research (IAAR), Chiba University, 1-33 Yayoi, Inage, Chiba, Chiba, 263-8522, Japan

**Keywords:** bud-sport, *Diospyros kaki* Thunb, transcriptome, tree crop

## Abstract

Fruit shape is an essential agronomic trait that contributes to the commercial value of fruit crops; however, the physiological mechanisms determining fruit shape have been studied only in a few model crops. Persimmon (*Diospyros kaki* Thunb.), a major fruit crop in East Asia, shows diversity in fruit shape among cultivars. The underlying molecular mechanisms that determine fruit shape remain unclear. A comparative physiological analysis was conducted to investigate the differences between a major Japanese cultivar, ‘Hiratanenashi’ with a “flat” fruit shape, and its somaclonal mutant cultivar, ‘Koushimaru’ with a “round” fruit shape. Through principal component analysis (PCA) with the elliptic Fourier descriptor, the fruit shape transitions in these two cultivars were measured over the fruit differentiation period (40-53 days after anthesis). Histological observation of the mesocarp cells in the fruit shape differentiation stage indicated that not only cell proliferation patterns but also multiple vectors involving cell shapes or sizes are possible determinants of differences in fruit shape. Transcriptome analysis at the fruit shape differentiation stage detected differentially expressed genes between the two cultivars. The hierarchical clustering with expression patterns throughout the fruit development/maturation stages defined four main clusters, including candidate genes determining fruit shapes. We focused on one of these genes, DKAch04a32073.t1, annotated as *WUSCHEL related homeobox 13* (*WOX13*), which has been reported to be a key factor in callus formation (or stem cell proliferation) and the reconnection of organs in *Arabidopsis thaliana*. Future functional analyses of the candidate genes would provide a better understanding of the mechanisms underlying persimmon fruit shape.

## Introduction

Plant organ shape has long been a subject of interest in the study of plant physiology and development, with the focus being mainly on the leaves (Tsukaya, 2005) or petals (Hileman, 2014). Fruit shape, although an important agronomical trait determining commercial value, has not been well investigated. In most studies involving fruit shape, the focus crop has been tomatoes (and potentially in the family Cucurbitaceae), and researchers have found that fruit shape may be explained by a small number of genes, such as *SUN, OVATE*, and *FAS1* (Wang et al., 2019). However, these genes cannot generally explain fruit shape in other crop species, including cucurbit species (Boualem et al., 2022; Xie et al., 2023), peach (Cirilli et al., 2021), and persimmon (Maeda et al., 2019). Little is known about the molecular mechanisms that determine fruit shape in tree crops, mainly because of the difficulty of genetic analysis, given the long juvenile phase or frequent polyploidy of these plants.

Persimmon (*Diospyros kaki* Thunb.), an important fruit tree crop that originated in East Asia, is cultivated mainly in China, Korea, and Japan. In terms of fruit shape, persimmons retain multidimensional diversity among cultivars. The fruit is not only differentiated as flat (e.g., ‘Hiratanenashi’, major cultivar in Japan) to narrow (e.g., ‘Saijo’, cultivated mainly in west Japan), but may also have grooves (e.g., ‘Yotsumizo’, a Japanese cultivar) or lid-like parts (e.g., ‘Tamopan’, Chinese cultivar). Although “shape” has been difficult to quantify, principal component analysis (PCA) with elliptic Fourier descriptors, along with analysis through SHAPE software (Iwata and Ukai, 2002) has enabled clustering of developing fruit shapes in a wide variety of persimmon cultivars (Maeda et al., 2018). Histological analysis of plants showing different fruit shapes (as characterized using SHAPE) suggests that differences in cell numbers or cell proliferation, rather than cell size or cell shape, determine persimmon fruit shape (Maeda et al., 2019). Numerical fruit shape development patterns among persimmon cultivars have also been investigated through co-expression network analysis, suggesting that persimmons have evolved a lineage-specific pathway to determine fruit shape (Maeda et al., 2019). This differs from the fruit shape-determining factors in tomatoes and applies the cytokinin-associated pathways represented by *knotted-like homeobox of Arabidopsis thaliana 1 (KNAT1)* (or genes involved in cell proliferation) to the shape-determining pathway (Maeda et al., 2019). Reports of synthetic cytokinin treatment affecting fruit shape (Itai et al., 1995) and significant differences in phytohormone levels between flat- and long-shaped persimmon fruits (Li et al., 2024) suggest that cytokinins and potential crosstalk between cytokinins and other phytohormones play important roles in persimmon fruit shape formation. Through population genetic approaches adjusted to polyploid species, researchers have identified several loci significantly associated with persimmon fruit shape, including variety-specific fruit shape-determining candidates (Horiuchi et al., 2023b). Tissue-specific transcriptome analysis of the side grooves of persimmon fruits suggests that *YABBY* homologs are most closely associated with groove depth (Kusumi et al., 2024). Based on these reports, it is likely that each lineage independently establishes a specific fruit shape. To comprehensively understand the diverse regulatory pathways underlying fruit shape determination, it is necessary not only to conduct extensive analyses across cultivars but also to focus on the molecular pathways specific to each lineage.

Tree crops often acquire new traits that arise from somatic mutations, a phenomenon known as “bud-sport”. A typical example of bud-sport is the derivation of early maturing varieties of Satsuma mandarin (*Citrus unshiu*) from a single genetically identical individual (Tanaka, 1925; Shimizu et al., 2017). Other examples of bud-sport in tree crops include skin color in oranges and grapes (Butelli et al., 2012; Walker et al., 2006), ripening timing in apples (Dong et al., 2011), fruit size variations in apples and European pears (Malladi and Hirst, 2010; Isuzugawa et al., 2014), and disease resistance in grapefruits (Ramekar et al., 2024). In case of bud-sport, it would be beneficial to conduct a comparative analysis between the derived and original accessions, with the goal of estimating a nearly identical genetic background and a single or limited number of variations (or polymorphisms). In the case of persimmon, a well-distributed bud-sport in Japan would be early-maturing derivatives from ‘Hiratanenashi’, such as ‘Tonewase’. A comparative transcriptome analysis between ‘Hiratanenashi’ and a derived smaller fruit cultivar, ‘Totsutanenashi’, revealed that the genes relating cell cycle were significantly down-regulated in ‘Totsutanenashi’ (Habu et al., 2016).

‘Koushimaru’, another example of a bud-sport cultivar derived from ‘Hiratanenashi’, was discovered in 1983 at Tsuruoka city, Yamagata prefecture, and officially registered in 1991. ‘Koushimaru’ produces more narrow or round fruits than does the original flat ‘Hiratanenashi’ with a pointed fruit apex (often described as round-shaped fruit when mature) (**Supplementary Fig. S1**). In this study, we conducted a comparative physiological analysis of ‘Hiratanenashi’ and ‘Koushimaru’. Based on chronological quantification of fruit shapes in these two cultivars by SHAPE, we aimed to identify the fruit shape differentiation stages, followed by histological and transcriptomic approaches in the fruit mesocarp to extract the candidate genes associated with the ‘Koushimaru’-specific fruit shape differentiation.

## Materials and Methods

### Plant materials

From May (stage immediately before anthesis) to September/October (fruit maturing stage) in 2020-2022, fruits were harvested approximately every 45 days, from a single identical tree of persimmon (*Diospyros kaki* Thunb.) cultivars, ‘Hiratanenashi’ and its somaclonal mutant ‘Koushimaru’ (**Supplementary Fig. S1**). Both cultivars were maintained at the Grape and Persimmon Research Station, NARO Institute of Fruit Tree and Tea Science (Hiroshima, Japan). The detailed sampling conditions for each experiment are described below.

### Fruit shape quantification through PCA with elliptic Fourier descriptors

Three to six fruits per experimental plot were sampled between May, September/October in 2020. Fruit shapes were quantified by PCA with elliptic Fourier descriptors, using SHAPE software (Iwata and Ukai, 2002), as described in a previous study (Maeda et al., 2018). After the calyx was removed from the fruit, it was cut longitudinally through the center. The fruit sections were photographed using a digital camera (TG-4, Olympus) at a distance of approximately 30 cm from the samples with a black background. An index scale of 3 cm × 3 cm and contour information were transformed into coefficients of elliptic Fourier descriptors as inputs in PCA using the SHAPE program. To identify the differences in the fruit shape differentiation stage between ‘Hiratanenashi’ and ‘Koushimaru’, the first principal component (PC1) value using all chronological samples throughout the fruit development/maturing stages were aligned to detect significant differences between the two cultivars, applying Student’s *t*-test in R function.

### Histological observation of mesocarp cells

Using fruits sampled in 2020, mesocarp sections at 40 days after anthesis (DAA) (at the end of June; Jun/E) and 65 DAA (at the end of July; Jul/E) were prepared with a cryostat (CM1520, Leica Biosystems), observed under an optical microscope, and photographed with a digital microscope camera (DP74-SET-A, Olympus). Two fruits were sampled per experimental plot, and 20-30 cell outlines, excluding tannin and vascular cells, were extracted from a section of each fruit. The cell areas were measured using ImageJ (Schneider, 2012) (50 cells per experimental plot). Cell shape was quantified using the SHAPE. Cell shape PC1 values were compared between ‘Hiratanenashi’ and ‘Koushimaru’ around the fruit shape differentiation stage.

### cDNA library preparation and Illumina sequencing

In order to identify differentially expressed genes (DEGs) between ‘Hiratanenashi’ and ‘Koushimaru’, 16 to 17 fruits of each cultivar were sampled on the early fruit shape-differentiating stage (June 13, 27 DAA), 2022. In the clustering analysis using time-series gene expression patterns of DEGs, two to four fruits per experimental plot were sampled at the flowering stage (May 19, 2 DAA), fruit development stage (June 29, 43 DAA; July 16, 60 DAA; August 2, 77 DAA), and mature stage (October 18, 124 DAA) in 2021. The RNA-seq library was prepared as previously described (Maeda et al., 2019). Briefly, total RNA was extracted from the mesocarp (10 mm × 5 mm × 3 mm) of fruit samples using the hot borate method (Wan and Wilkins, 1994). mRNA was isolated using the Dynabeads™ mRNA purification kit (Ambion) from 400-500 ng of total RNA. cDNA libraries were prepared using the KAPA RNA HyperPrep Kit (Roche) and purified using AMPureXP (Beckman Coulter Life Sciences) to remove short fragments. The libraries were sequenced on Illumina Hiseq 4000 platform at the QB3 Genomic Sequencing Laboratory (or Vincent J. Coates Genomics Sequencing Laboratory) of UC Berkeley (http://qb3.berkeley.edu/gsl). Raw sequencing data were processed using custom Python scripts (https://github.com/Comai-Lab/allprep/blob/master/allprep-13.py) in order to de-multiplex and trim the read data, as previously described (Akagi et al., 2014).

### Transcriptomic analysis

The mRNA-seq reads were aligned to the reference CDS sequences of *Diospyros kaki* Thunb. ‘Taishu’ (Horiuchi et al., 2023a) (https://persimmon.kazusa.or.jp/index.html) using the Burrows-Wheeler aligner (BWA) (version 0.7.15) (Li and Durbin, 2009) with default parameters, and converted to SAM file using SAMtools (Li and Durbin, 2009). Mapped reads were counted per CDS using the custom R program to calculate read number per kilobase and mapped million reads (RPKM) as normalized gene expression. PCA analysis was conducted with the RPKM values (only > 1), using the prcomp function in R. DEGs were detected during fruit shape differentiation stage between ‘Hiratanenashi’ and ‘Koushimaru’ sampled in 27 DAA of 2022, using DESeq2 (Love et al., 2014) (FDR < 0.01: DESeq2, *P* < 0.1: Student’s *t*-test). The TAIR10 database (https://www.arabidopsis.org/index.jsp) was used for functional annotation of the DEGs. Using fruits chronologically sampled in 2021, hierarchical clustering for gene expression patterns of DEGs were processed using hclust function in R. GO enrichment analysis was conducted with ShinyGO 0.77 (Ge et al., 2020) (http://bioinformatics.sdstate.edu/go/). The threshold for the significance of enriched GO terms was set at FDR < 0.05.

## Results and Discussion

### Transition of fruit shapes

The numerical fruit shape characterization using SHAPE program indicated that the PC1 accounted for 68.19 % of the total explanatory variables and mainly reflected the continuous changes of flatness-roundness between ‘Hiratanenashi’ and ‘Koushimaru,’ whereas PC2 accounted for 10.70 % of the total explanatory variables and mainly reflected the growth stage (**Fig. 1A**). The distribution of PC1 and PC2 values over time indicated that biological replicates exhibited a large degree of variation within a sample in the early developmental stages, whereas they converged into distinct trends specific to each cultivar in later developmental stages (**Supplementary Fig. S2**). No significant difference was detected in the PC1 value between the two cultivars until 40 DAA (at the end of June; Jun/E). After 53 DAA (at the beginning of July; Jul/B), the PC1 values differed at all sampling points (*P* = 0.002, **Fig. 1B**). Thus, we defined that the duration from 40 DAA to 53 DAA as the fruit shape differentiation stage between ‘Hiratanenashi’ and ‘Koushimaru’.

**Fig. 1.**
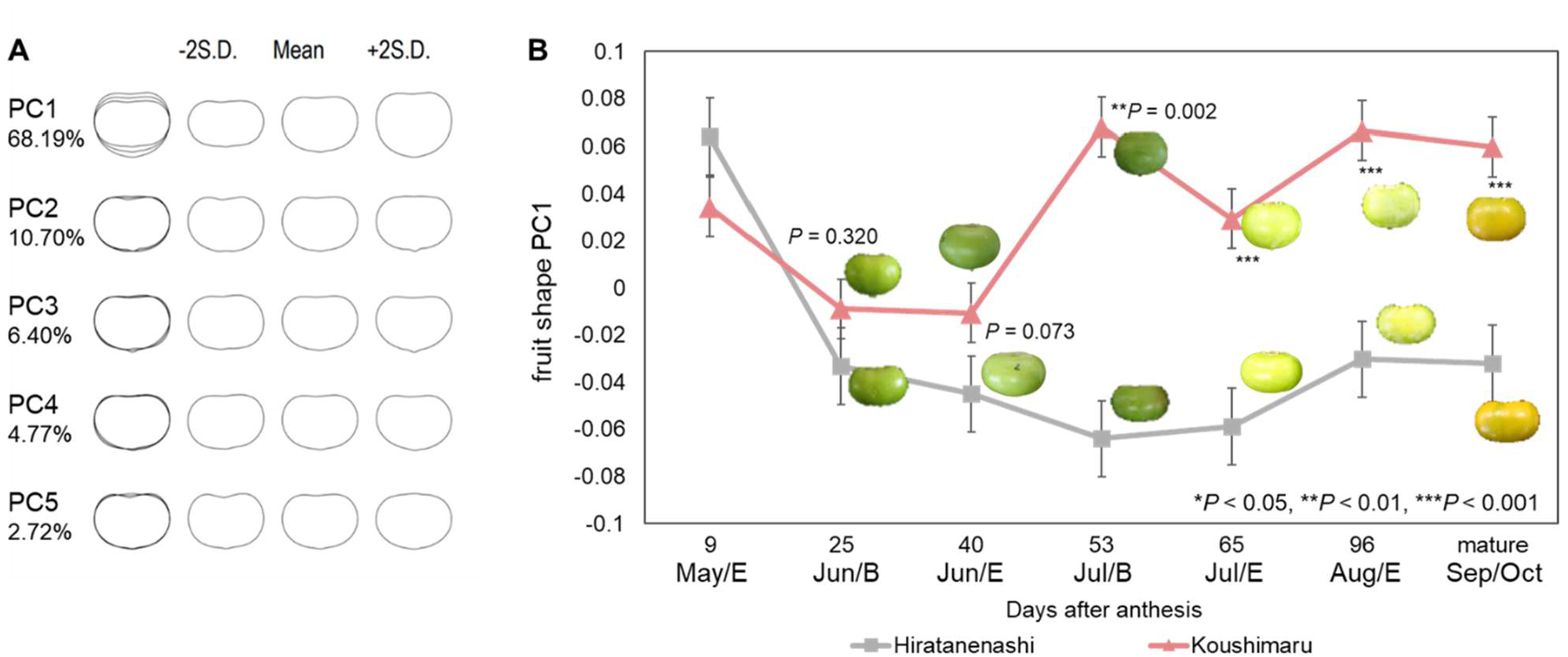
Fruit shape quantification and transition from the fruit development stage to the mature stage between ‘Hiratanenashi’ and its somaclonal mutant cultivar ‘Koushimaru’. SHAPE analysis was performed using longitudinal fruit sections (ovary sections in anthesis) of ‘Hiratanenashi’ and ‘Koushimaru’ from May to September/October (mature stage). The contribution value of each PC is expressed as a percentage. **(A)** The longitudinal fruit shape was reconstructed using an elliptical Fourier based on the PC1-PC5 axes. **(B)** Comparison of the fruit shape PC1 characterized by SHAPE analysis from the fruit development stage to the mature stage between ‘Hiratanenashi’ and ‘Koushimaru’. “month/B”, “month/M”, and “month/E” mean the beginning of month, the middle of month, and the end of month, respectively. Photographs of the average longitudinal sections of the fruit at each growth stage are shown.

### Cell division and elongation patterns in the fruit shape differentiation stage

To investigate the histological difference between ‘Hiratanenashi’ and ‘Koushimaru’, cell shape and size were observed from mesocarp sections under microscopy, in the fruit shape differentiation stage (**Fig. 2A**). The results of the SHAPE program indicated that cell shape PC1, which accounted for 47.80 % of the total explanatory varieties, mainly reflected changes in the cell aspect ratio (**Supplementary Fig. S3**), which was consistent with the analysis of the longitudinal section of the fruit (**Fig. 1A**). ‘Koushimaru’ showed significantly higher PC1 values than did ‘Hiratanenashi’ at 40 DAA (*P* = 0.040, **Fig. 2B**). Although there were no significant differences in fruit shape between two cultivars at 40DAA (**Fig. 1B**), differences in cell shape at this stage may contribute to later differences in fruit shape. The results of cell area measurements by ImageJ (Schneider, 2012) between ‘Hiratanenashi’ and ‘Koushimaru,’ showed that the cell area of ‘Hiratanenashi’ was significantly larger than that of ‘Koushimaru’ at 65 DAA (*P* < 0.010, **Fig. 2C**). The results of previous studies of various persimmon cultivars have suggested that the diversity of longitudinal fruit shape is determined by cell proliferation levels rather than cell shape or elongation patterns (Maeda et al., 2018, 2019). On the other hand, our findings suggest that not only cell proliferation but also cell shaping and/or sizing would be key factors to contributing to persimmon fruit shape, at least in comparison of ‘Hiratanenashi’ and ‘Koushimaru’.

**Fig. 2.**
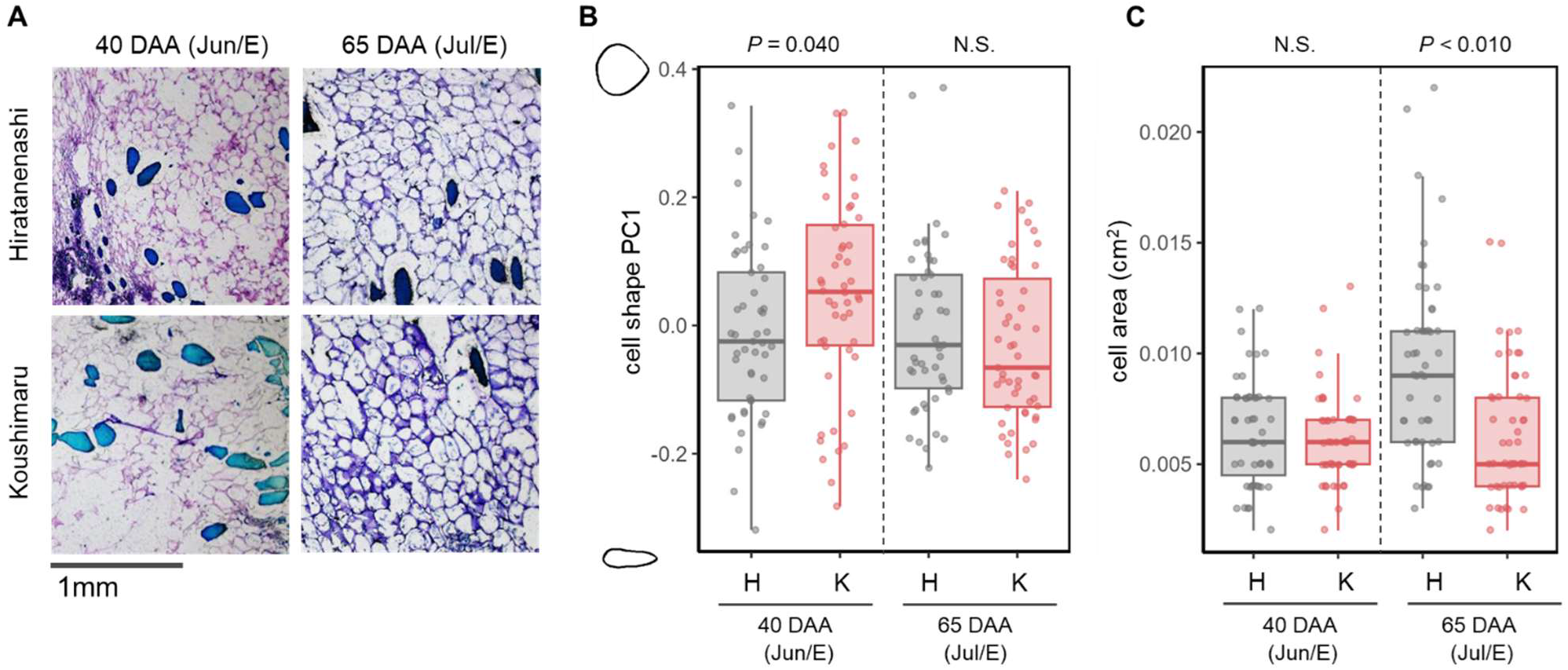
Observation of cell division and elongation process in the fruit differentiation stage. **(A)** Tissue section of the mesocarp at 40 and 65 days after anthesis (DAA). The sliced mesocarps were dyed with the Toluidine blue. The scale bar length is 1 mm. Darkly stained areas are the tannin cells. **(B)** Comparison of the cell shape PC1 characterized by SHAPE analysis at 40 and 65 DAA. ‘Hiratanenashi’ is plotted in gray boxes and ‘Koushimaru’ is plotted in pink boxes. “month/E” means “the end of month”. **(C)** Comparison of the cell area measured by ImageJ at 40 and 65 DAA. ‘Hiratanenashi’ is plotted in gray boxes and ‘Koushimaru’ is plotted in pink boxes. “month/E” means “the end of month”.

### Transcriptome analysis in the fruit shape differentiation stage

Transcriptome analysis of the mesocarp in the fruit shape differentiation stage detected 822 DEGs between ‘Hiratanenashi’ and’ Koushimaru’ (**Fig. 3A**; FDR < 0.01: DESeq2, *P* < 0.1: Student’s *t*-test). The DEGs consisted of 594 up-regulated and 228 down-regulated in ‘Koushimaru’, in comparison to ‘Hiratanenashi’. GO enrichment analyses on them, using the GO Biological Process (BP) pathway database, detected the terms involving response to ethylene and other hormone (marked with a red circle) in the DEGs up-regulated in ‘Koushimaru’ (**Fig. 3B**), and the terms involving response to low oxygen condition (or hypoxia) in the DEGs both up-regulated and down-regulated in ‘Koushimaru’ (**Fig. 3B, C**, marked with a black circle). These trends are similar to what occurs with auxin signal transduction (Eysholdt-Derzsó and Sauter, 2017; Hartman et al., 2019, 2021; Shukla et al., 2019). The network between GO terms further suggested a functional link between phytohormone signaling pathways and those involved in responses to hypoxic environments (**Supplementary Fig. S4**). Consistent with this situation, some GOs associated with the phosphorylation relay (**Fig. 3B, C**, marked with a green circle), as the main signal transduction driver, were enriched. In summary, GO enrichment analysis suggested that, especially in ‘Koushimaru’, ethylene- or auxin-related signaling may be highly active.

**Fig. 3.**
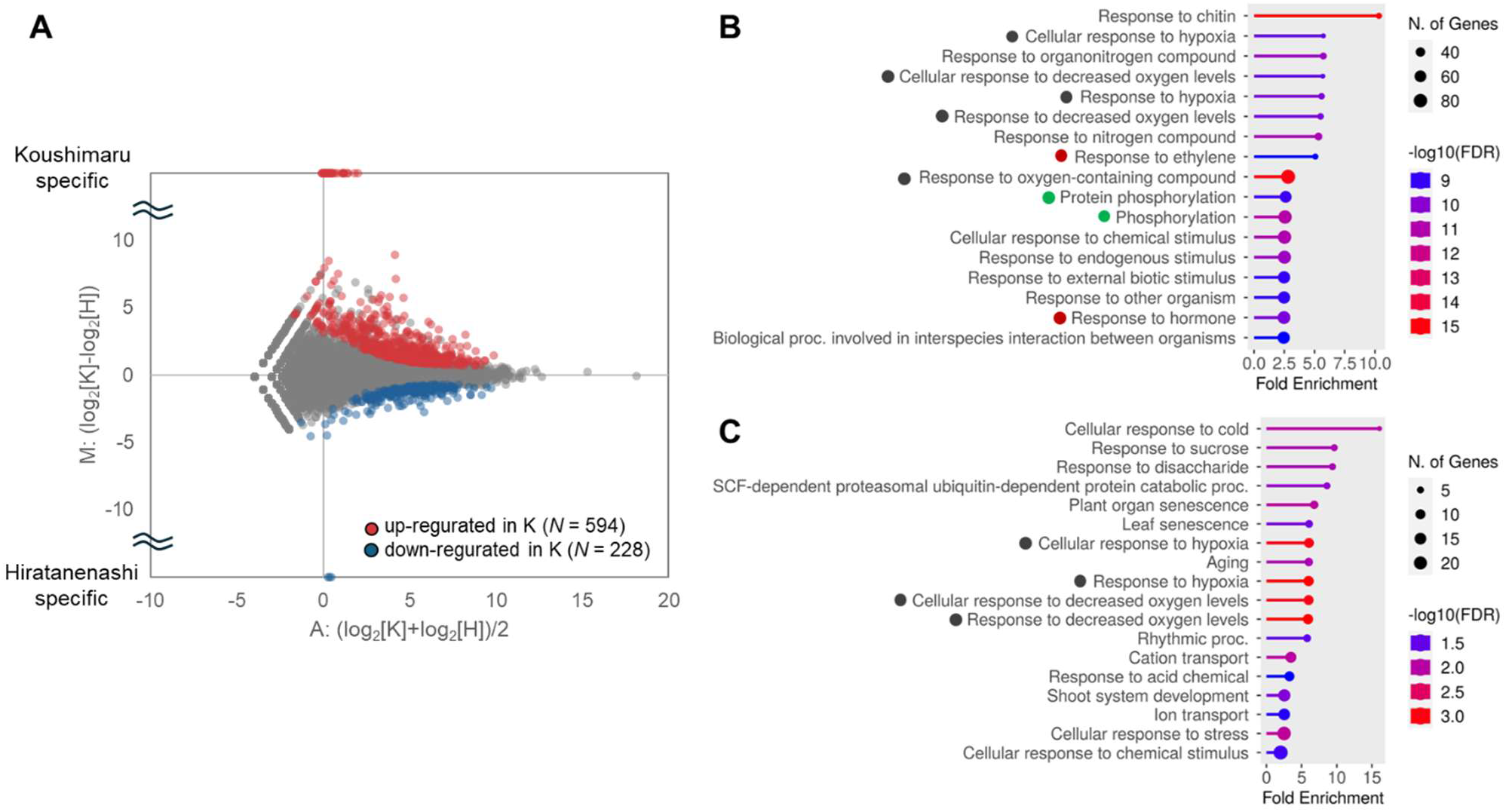
Comparative transcriptome analysis between ‘Hiratanenashi’ and ‘Koushimaru’. **(A)** M-A plot of the transcriptomic data of ‘Hiratanenashi’ and ‘Koushimaru’ at 27 DAA. Differentially expressed genes (DEGs) were colored with red (up-regulated in ‘Koushimaru’) or blue (down-regulated in ‘Koushimaru’). **(B)** Top GO terms enriched among DEGs up-regulated in ‘Koushimaru’. Gene ontology (GO) terms associated with hypoxia are marked with a black circle, those related to hormone are with a red circle, those associated with phosphorylation with a green circle. **(C)** Top GO terms enriched among DEGs down-regulated in ‘Koushimaru’. GO terms associated with hypoxia are marked with a black circle.

Hierarchical clustering with chronological expression transitions from May to September/October (mature stage) (in 2021) categorized 753 of the 822 DEGs into 13 clusters (**Fig. 4A**, height = 7.5, Hclust). The other 69 DEGs did not satisfy the quality or read counts required to properly conduct clustering with the chronological transcriptomic data in 2021. Clusters were integrated based on their stage-specific expression trends, and the DEGs were classified into four groups. Group 2, which comprised 179 DEGs, exhibited expression changes (including highly/low expression) specific to the fruit shape differentiation stage (or the end of June (Jun/E)). These genes had significantly enriched GO terms related to phytohormones (**Fig. 4B**; marked with a red circle), which included genes annotated to abscisic acid, salicylic acid, jasmonic acid, and ethylene transduction pathways, all of which are known to be involved in senescence and in stress responses (Horváth et al., 2007; Iqbal et al., 2017; Tuteja, 2007; Wang et al., 2020). Genes related to the auxin, cytokinin, and brassinosteroid pathways, which play key roles in plant growth and development (Müssig, 2005; Werner et al., 2001; Woodward, 2005), were also included. These results suggest that a complex regulatory mechanism involving crosstalk among multiple phytohormone pathways may explain the observed differences in fruit shape between the two cultivars.

**Fig. 4.**
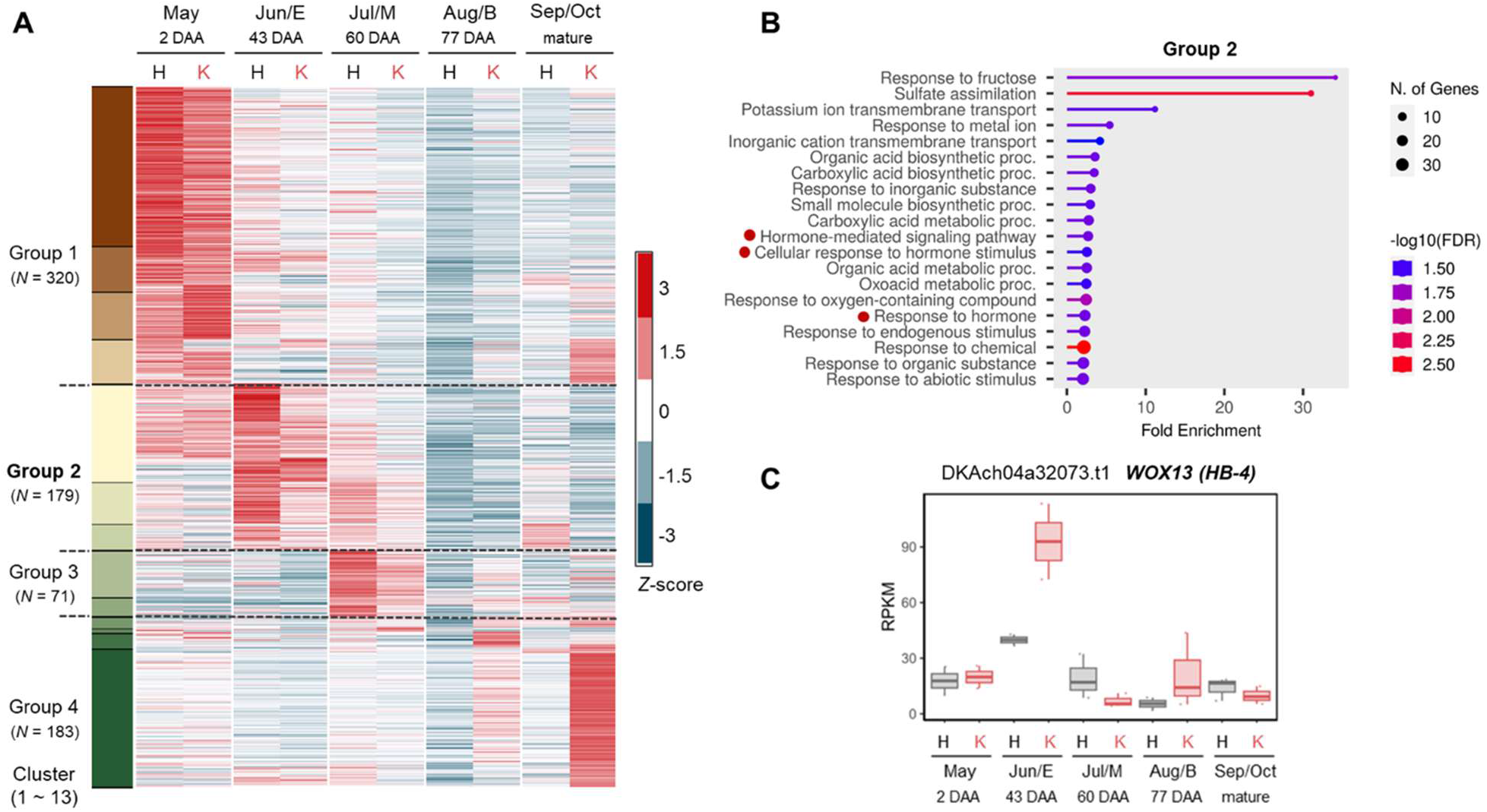
Clustering of DEGs using transcriptomic data over fruit development stage. **(A)** The heatmap shows that the results of hierarchical clustering based on gene expression patterns from May to September/October of DEGs. “month/B”, “month/M”, and “month/E” mean the beginning of month, the middle of month, and the end of month, respectively. “H” and “K” means ‘Hiratanenashi’ and ‘Koushimaru’ respectively. **(B)** The list of top GO terms obtained by GO enrichment analysis for DEGs in Group 2. GO terms associated with hormones are with a red circle. **(C)** Gene expression patterns from May to September/October (mature stage) of DKAch04a32073.t1 in Group 2. ‘Hiratanenashi’ is plotted in gray boxes and ‘Koushimaru’ is plotted in pink boxes. “month/B”, “month/M”, and “month/E” mean the beginning of month, the middle of month, and the end of month, respectively.

Here, we focused on DKAch04a32073.t1 in Group 2, which is annotated as *WUSCHEL related homeobox 13* (*WOX13*) belonging to the WOX transcription factor family, and more highly expressed in the somaclonal mutant ‘Koushimaru’ than in ‘Hiratanenashi’ at Jun/E (**Fig. 4C**). Recently, *WOX13* was reported to be a key player in controlling callus formation and organ reconnection in *Arabidopsis thaliana* (Ikeuchi et al., 2022). Thus, *WOX13* can inactivate cell differentiation and stem cell proliferation through the reception of wounding signals. Consistent with this concept, auxin-induced *WOX13* negatively regulates the shoot apical meristem (SAM) formation and promotes callus formation by stimulating the expression of *EXPA17* in *A. thaliana* (Ogura et al., 2023). There have also been reports that *WOX13* is expressed in the replum, which is the silique of *A. thaliana*, and promotes replum development by preventing the activity of the *JAG* (*JAGGED*, C2H2 Zn-Finger encoding gene) */FIL* (*FILAMENTOUS FLOWER, YABBY1* (*YAB1*) group genes) (Romera-Branchat et al., 2013). In tree crops, it has been reported that *VviWOX13C* plays a key role in fruit set; for example, in *Vitis vinifera* (Sun et al., 2024). Although these reports do not directly support the involvement of *WOX13* in fruit shape determination, the suggested pathways may be associated with mesocarp cell proliferation under stressful conditions. Namely, based on our comparative transcriptome analysis, higher expression of DKAch04a32073.t1 in ‘Koushimaru’ may inhibit cell differentiation (or activate stem cell proliferation) under ethylene/auxin (or its related) signals in mesocarp tissue, potentially resulting in fruit shape differentiation between the two cultivars. Consistent with *WOX13* regulation in *Arabidopsis*, the enriched GO terms from the DEGs suggested the involvement of auxin signal transduction. DKAch06a20756.t1 and DKAch07a36577.t1, annotated as the auxin-inducible genes *IAA5* and *IAA14*, respectively, and DKAch03a26128.t1, annotated as *EXPA15* were also detected in Group 2. They would further support the possibility that DKAch04a32073.t1 (or *WOX13*) induced by auxin signals regulates *EXPA* gene(s), to determine the fruit shape in ‘Koushimaru’. The involvement of the cytokinin signaling pathway in fruit development and fruit shape determination in persimmons has been widely reported (Itai et al., 1995; Maeda et al., 2019; Sobajima et al., 1974), whereas auxin-related genes identified in gene network analyses are also associated with persimmon fruit shape (Kusumi et al., 2024). Cytokinin and auxin signaling pathways are known to often function antagonistically (Moubayidin et al., 2009); however, the crosstalk between them in fruit shape regulation in persimmons remains unclear. To clarify these mechanisms, it is necessary to demonstrate the function of the described candidate genes, including DKAch04a32073.t1, and to investigate the hierarchical relationships between them.

## Conclusion

We conducted a comparative analysis using persimmon cultivar ‘Hiratanenashi’ and its somaclonal mutant cultivar ‘Koushimaru’. As a result, we defined 40–53 DAA as the fruit shape differentiation stage between the two cultivars, which approximately coincides with the stage defined in the analysis of many persimmon varieties in a previous study (Maeda et al., 2018). Morphological observations of the mesocarp cells in the fruit shape determination stage suggested that there might be differences not only in the direction of cell proliferation (Maeda et al., 2018) but also in cell shape and size between the two cultivars. Comparative transcriptome analysis at the fruit differentiation stage allowed us to efficiently identify a group of genes that are important for determining fruit shape. From this analysis, we could extract new candidate genes, such as DKAch04a32073.t1 annotated as *WUSCHEL related homeobox 13* (*WOX13*), which may be related to the difference in fruit shape between ‘Hiratanenashi’ and ‘Koushimaru’. These findings enhance our understanding of the genetic pathways involved in fruit shape determination in persimmons. Further research is necessary to determine the biological and molecular functions of these candidate genes.

## Acknowledgments

We are grateful to all members of the NARO Institute of Fruit Tree and Tea Science for maintaining the genomic resources of Oriental persimmon. This work was supported by the JSPS Grants-in-Aid for Scientific Research, grant numbers 23K23604, 24KJ1720, and 25K00086.

## Author contribution statement

A.H. and T.A. conceived the study and designed experiments. A.H. conducted the experiments. A.H. and T.A. analyzed the data. R.M., N.O., K.U., Y.K. and T.A. contributed to plant resources and facilities. A.H., T.A. and M.F.M. drafted the manuscript. All authors approved the manuscript.

