## Supplemental Figures for "Physiological characterization in a somaclonal mutant for fruit shape derived from persimmon ‘Hiratanenashi’"

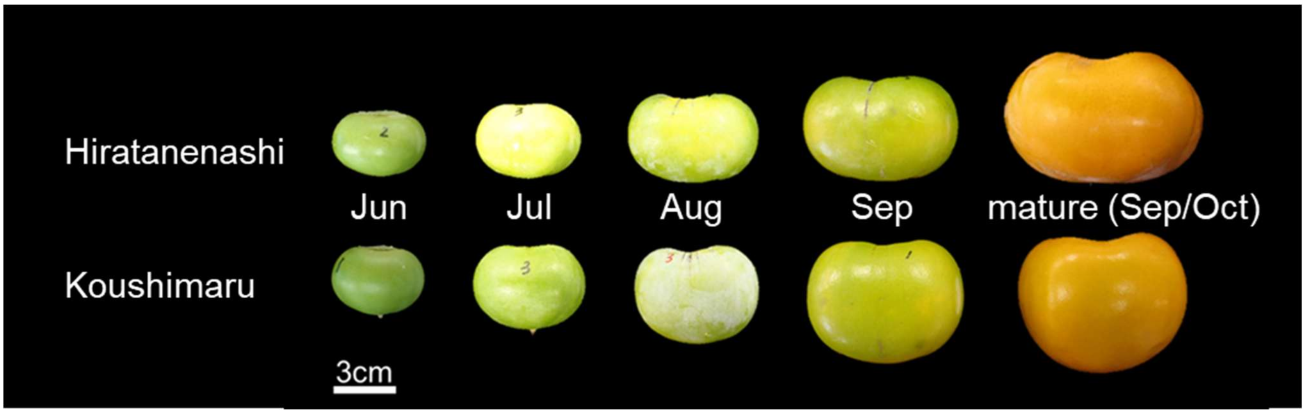


**Supplementary Fig. S1**

Photographs in the longitudinal direction of ‘Hiratanenashi’ and ‘Koushimaru’ fruits from the fruit developmental stage to the mature stage. Scale bar, 3 cm.

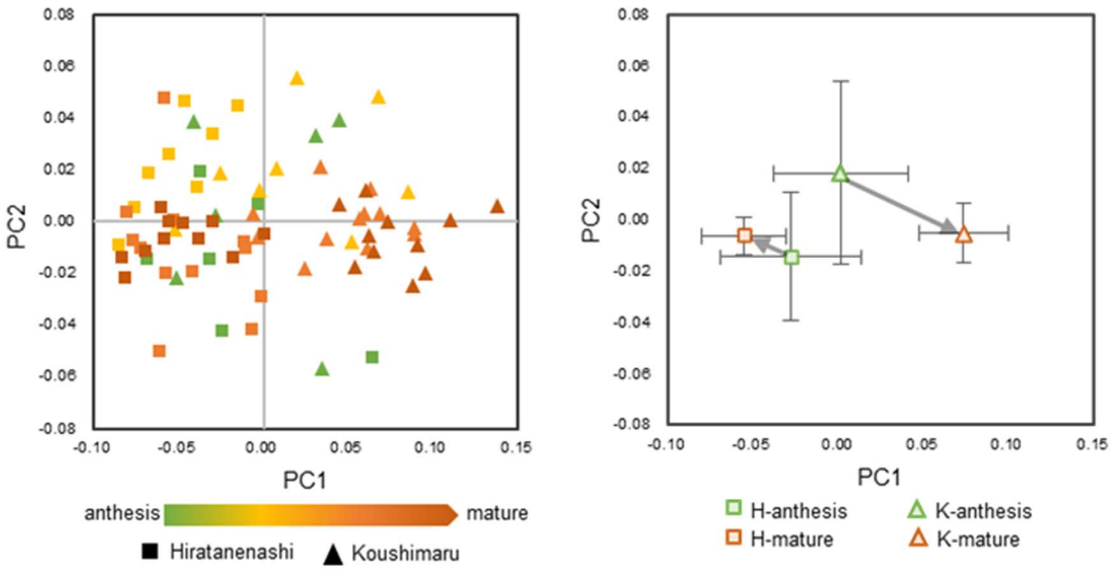


**Supplementary Fig. S2**

SHAPE analysis using the longitudinal fruit sections (use ovary sections in anthesis) of 'Hiratanenashi' and 'Koushimaru' from May to September/October (mature stage). The contribution value of each PC is shown as a percentage. The left dot plot shows the distribution of fruit shape PC1 and PC2 values. The dots were colored with growth stages. The right plot shows the transition of fruit shape PC1 and PC2 values from anthesis to mature stage.

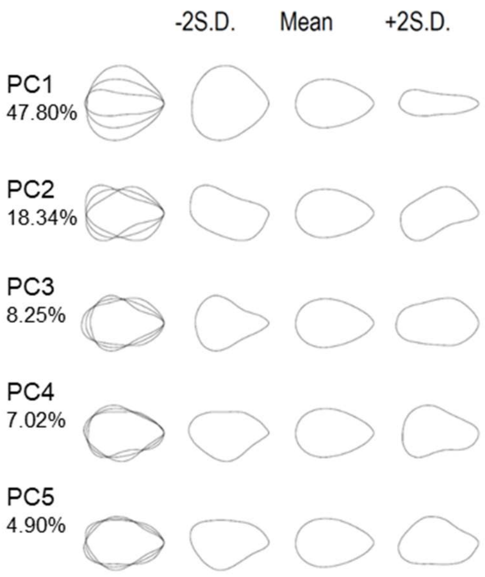


**Supplementary Fig. S3**

SHAPE analysis using the cell outlines of ‘Hiratanenashi’ and ‘Koushimaru’ at 40 and 65 DAA. The contribution value of each PC is shown as a percentage. Fruit longitudinal shape was reconstructed from coefficients of elliptic Fourier descriptors for PC1-PC5 axes.

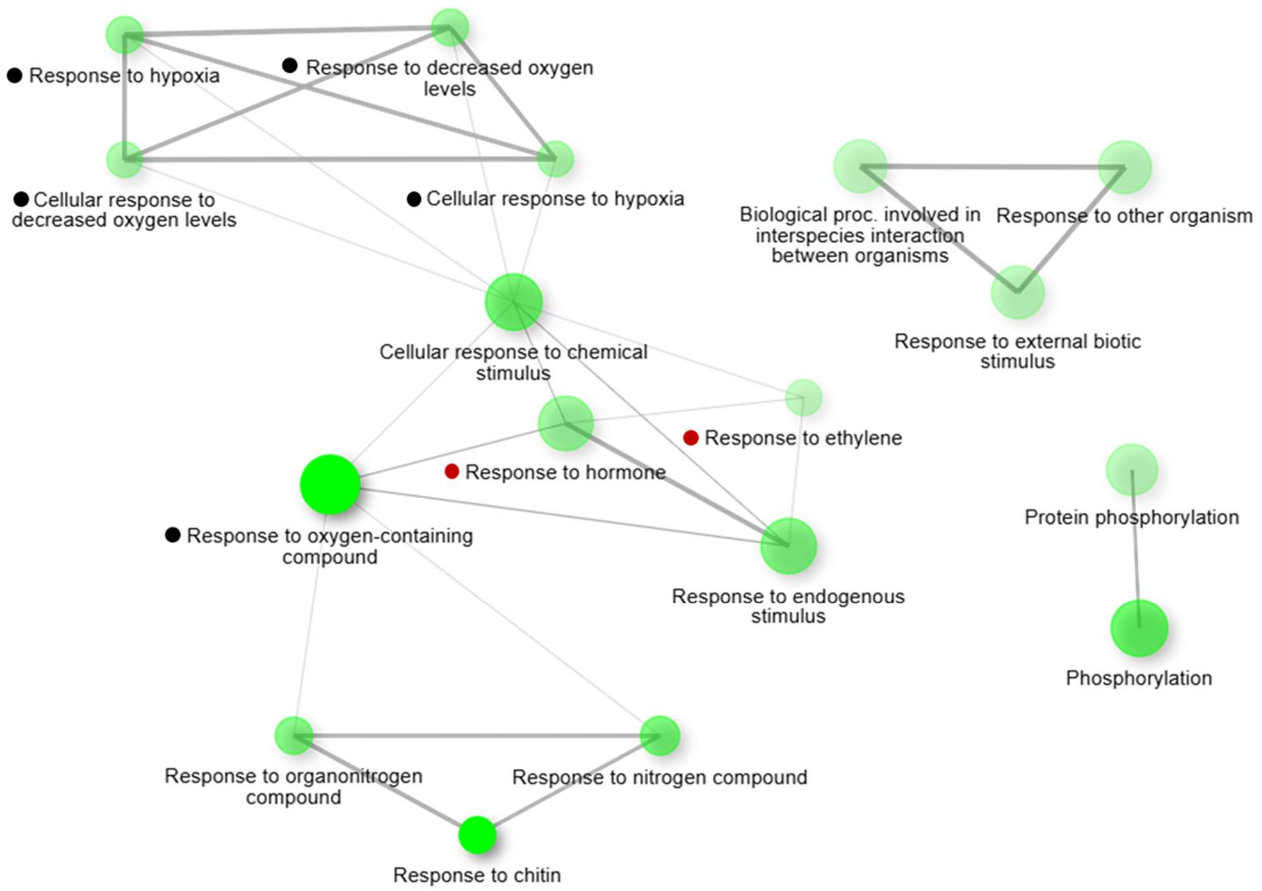


**Supplementary Fig. S4**

Interactive plot of the relationship between enriched pathways among DEGs up-regulated in 'Koushimaru'. Darker nodes are more significantly enriched gene sets. Larger nodes represent larger gene sets. Thicker edges represent more overlapped genes. Black circle, GO terms associated with hypoxia; red circle, terms related to hormones.
